# Genome-wide identification and analysis of analysis of BURP genes in *Triticum aestivum*

**DOI:** 10.1101/2022.10.21.513156

**Authors:** Wengen Zhu

## Abstract

Plant-specific BURP genes are involved in most aspects of plant development and in diverse stress responses. However, there has been no comprehensive and systematic research on the wheat (*Triticum aestivum* L.) BURP gene family. We comprehensively identified 54 BURP genes in wheat and 12, 23, and 32 BURP genes, respectively, in its three progenitor species, *Triticum urartu, Aegilops tauschii,* and *Triticum dicoccoides*. By phylogenetic analysis, we classified the wheat BURP genes into five subfamilies: BURP VI, BURP VII, RD22-like, PG1β-like, and the newly defined BURP VIII. BURP genes were distributed unevenly on 20 chromosomes, and 33 (61%) were anchored in the distal chromosome segments. Analysis of gene duplication events showed that segmental duplication was the main contributor to the expansion of this gene family in wheat. Assessment of tissue-specific and stress-induced expression indicated that most BURP members are heavily involved in plant development and responses to various stress conditions. RNA-seq data revealed ten *TaBURP* genes expressed specifically in spikes.

## Background

Wheat is one of the most important food crops worldwide, providing approximately 20% of energy for human consumption. Hexaploid wheat (*Triticum aestivum; 2n* = *6x* = 42; AABBDD) was derived from two hybridizations between three gramineous ancestors. The most recent hybridization event, which occurred around 8,000 years ago, was between cultivated tetraploid emmer wheat (*Triticum dicoccoides,* AABB) and diploid goat grass (*Aegilops tauschii,* DD). However, because of the enormous genome and complex structural features of wheat, investigating genetic function in this species is challenging. Fortunately, with the advancement of sequencing technology, high-quality genome assembly and annotation of wheat has been achieved, making detailed phylogenetic and functional characterization of gene family feasible in this species.

The BURP protein family is ubiquitous across plants but is not found in other organisms. BURP proteins, characterized by the possession of BURP domains, can be broadly classified into several subfamilies, including BNM2 (a microspore protein in Rape *[Brassica napus]*) [1], VfUSP (an unknown seed protein in field bean *[Vicia faba]*) [2], AtRD22 (a dehydration-responsive protein in Arabidopsis *[Arabidopsis thaliana]*) [3], LePG1β (a non-catalytic β-subunit of the polygalacturonase isozyme 1 in tomato *[Lycopersicon esculentum]*) [4] and others [5–7].

BURP-domain-containing proteins are generally conserved and comprise three or four modules. Specifically, the N-terminal hydrophobic domain is a potential signal peptide; the variable middle regions contain an optional short conserved segment or other short segments or a selectable segment composed of repetitive units; and the C-terminus is the conserved BURP domain, typically with a few conserved amino acid sites and four repetitive cysteine histidine (CH) motifs, also known as CHX_10_CHX_23-37_CHX_23-26_CHX_8_W (where X = any amino acid residue) [6, 8]. It is widely acknowledged that differences in the middle region are what make individual BURP proteins unique.

BURP genes have been phylogenetically and functionally described in diverse plants, such as cotton (*Gossypium hirsutum*) [6], soybean (*Glycine max*) [9], alfalfa (*Medicago sativa*) [10], common bean (*Phaseolus vulgaris*) [11], *Medicago truncatula* [12], rice (*Oryza sativa*) [5], maize (*Zea mays*), and sorghum (*Sorghum bicolor*) [7]. Their functions can be broadly categorized into two groups: those involved in plant development and those involved in responses to environmental stresses.

VfUSP provided early evidence for the role of BURP protein in regulating plant development. *VfUSP* is highly expressed in developing seeds of *V. faba* [2]. Ectopic expression of the Arabidopsis gene *AtUSPL1* disrupts seed development and modifies the morphology of seed lipid vesicles [13]. In *B. napus, BNM2* is expressed in microspore-derived embryos, and the protein is located in seed protein storage vacuoles [2, 14], whereas *BNM1* is specifically expressed in pollen [15]. *ASG1* is specifically expressed in apomixis-specific cells, indicating its potential function in embryo development [16, 17]. In rice, OsRAFTIN is specifically targeted to microspore exine and accumulates in Ubisch bodies. Suppression of *OsRAFTIN* expression leads to functional deficiency in the anther and affects anther development [18]. *RA8* is preferentially expressed in the rice anther tapetum, middle layer, and endothecium tissues, and its high expression might play a crucial role in the regulation of anther development [19].

*SCB1,* a BURP gene in soybean, is expressed in thin- and thick-walled parenchyma and may help control seed coat parenchyma cell differentiation [20]. AtUSP1 exists in cellular compartments and contributes critically to the production and storage of seed proteins [13]. In tomato, PG1β is a key localizer and regulator of pectin solubilization and degradation during fruit ripening [4, 21]. Thus, BURP genes are involved in most aspects of plant development.

Another pivotal function of BURP genes is response to diverse stresses. Some BURP gene subfamilies, especially the RD22-like and BNM2-like subfamilies, are hypersensitive to environmental stress [3, 22]. For example, two loss-of-function mutants, *rd22* and *atusp1,* in Arabidopsis show enhanced water stress tolerance [23]. The expression of *BnBDC1,* a homolog of *RD22* in *B. napus,* was up-regulated under osmotic stress and down-regulated after UV irradiation and salicylic acid (SA) treatment [24]. Toshiaki *et al*. observed differential expression of four genes homologous to *RD22* in mangrove (*Bruguiera gymnorrhiza*) under different stress conditions [25]. SBIP-355, a BURP gene homologous to RD22 in tobacco (*Nicotiana benthamiana),* might be involved in plant defense through the SA pathway [26]. Two soybean genes, *Sali5-4a* and *Sali3-2,* were induced by aluminum stress [27].

To date, numerous systematic genome-wide analyses of the BURP gene family have shown that most members participate in multiple stress responses [5, 7, 9, 10, 12], including abiotic stresses, and possibly in hormone signaling pathways such as the abscisic acid (ABA) and SA pathways [6]. These functions may facilitate plant adaptation to harsh environments. Despite the important of these proteins, there are no available studies on the wheat BURP gene family. Moreover, little is known about the relationship between wheat BURP genes and development. To fill these gaps, we comprehensively analyzed putative BURP genes in *T. aestivum* (AABBDD), as well as its three progenitor species: *Triticum urartu* (AA), *Aegilops. tauschii* (DD), and *Triticum. dicoccoides* (AABB). We provide a detailed overview of the members and their phylogeny. Furthermore, we identified tissue-specific expression of BURP genes from a public transcriptional database. We also investigated *TaBURP* expression in response to drought stress, salinity stress, cold stress, and ABA treatment.

## Results

### Identification and characterization of *BURP*

We identified a total of 54, 12, 23, and 32 putative BURP domain proteins in *T. aestivum, T. urartu, Ae. tauschii* and *T. dicoccoides,* respectively (. The 54 *TaBURP* genes were named on the basis of previous studies [28]. We assigned a uniform nomenclature to all wheat BURP genes based principally on their subfamily relationships, subgenomic positions (A, B, or D) and phylogenetic relationships, as previously described [28]. Specifically, individual gene names begin with the abbreviation of the species nomenclature (e.g., Ta for *Triticum aestivum*). Subsequently, genes were named based on previous classification of rice gene subfamily [5]. For example, ‘BURPVII’ represents the BURP VII subfamily. The only exceptions are the BURP VIII subfamily, which are not found in rice. Capital letters ‘A’, ‘B’, or ‘D’ indicate the subgenomic position of the gene, such as *TaRD22-A1*, *TaRD22-B1*, and *TaRD22-D1*. Genes originating from one subfamily but with distinct homeologous traits within the same genome were sequentially numbered: for example, *TaBURPVI-A2* and *TaBURPVI-A3*. Other putative homologs due to tandem duplication, transposition, or other reasons were named using successive numbers following a dash: for example, *TaBURPVII-B1-1* and *TaBURPVII-B1-2*). The length of TaBURP proteins ranged from 161 aa to 754 aa. The predicted molecular weight (Mw), isoelectric point (pI), grand average (GRAVY) of hydropathicity, and predicted subcellular localization of all wheat BURP domain proteins are shown in Table S1.

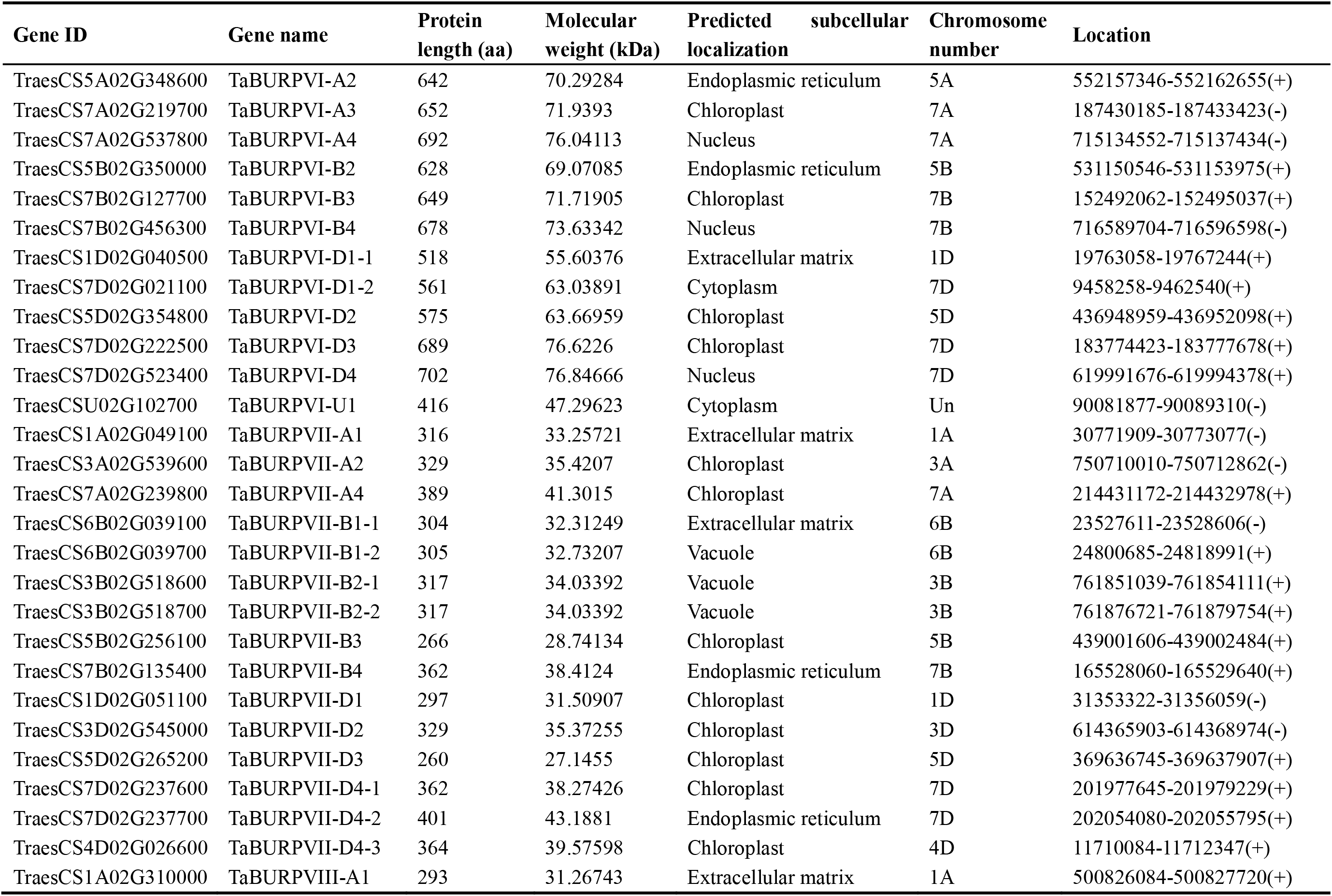
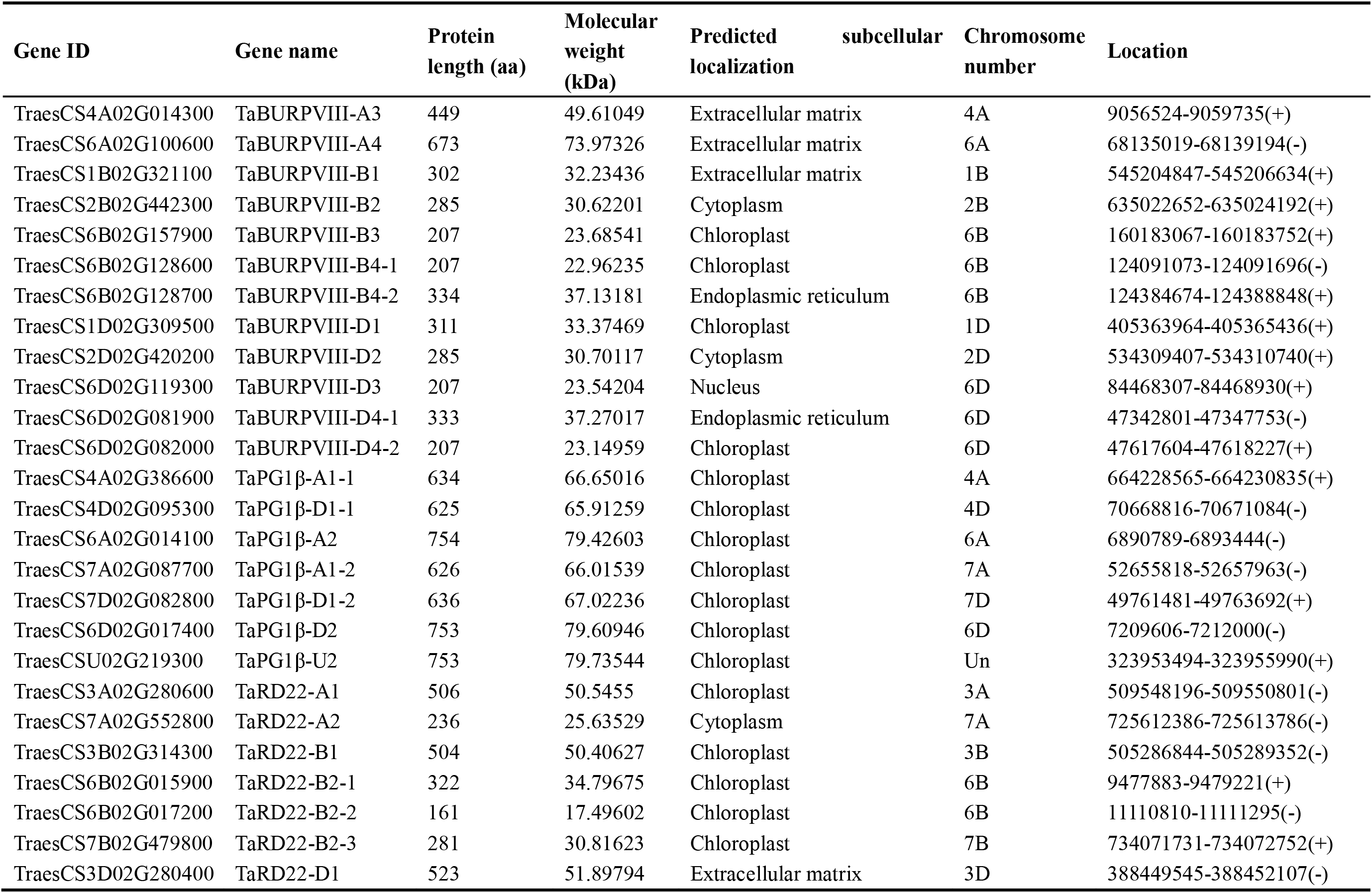

### Phylogenetic and classification analysis of BURPs

To investigate the evolutionary relationship of *TaBURPs* between wheat (54), rice (17), maize (10), Arabidopsis (5), soybean (26), and the four canonical proteins(BNM2, VfUSP, RD22, and LePG1β), a phylogenetic tree was inferred with maximum likelihood (Fig. 1, **Additional file 1: Table S3 and S4**). The differences in the numbers of BURP members in rice, maize, and soybean found here compared with previous reports may be associated with our selection of only one representative gene transcript and the selection of different genomic versions [6]. The phylogenetic tree showed that wheat BURP genes were clustered into five BURP subfamilies, including five well-defined subfamilies (BURP VI, BURP VII, BURP VIII, RD22-like and PG1 β-like).

**Fig. 1.**
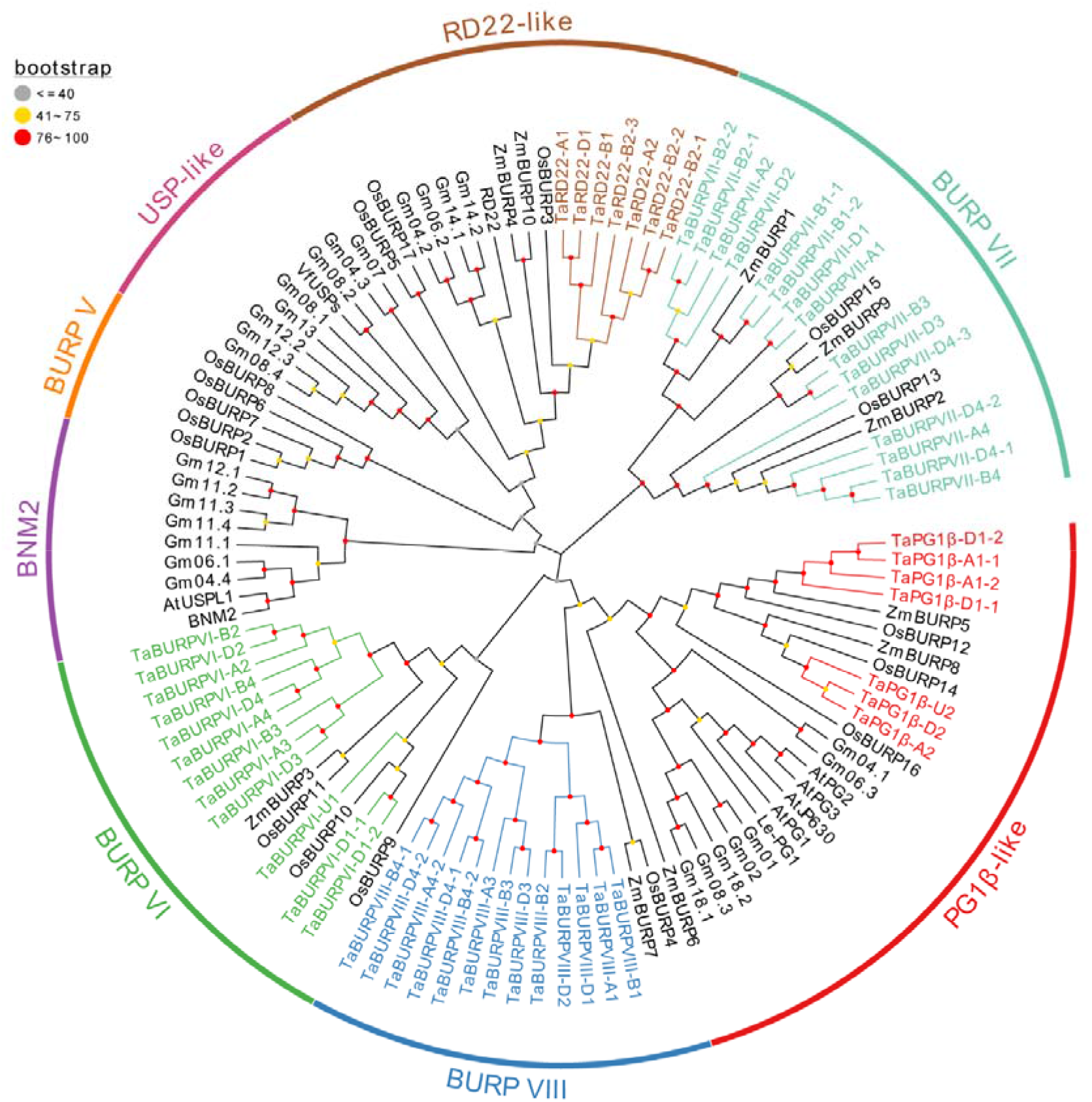
Phylogenetic tree of BURP proteins from wheat, Arabidopsis, soybean, maize, and rice. MEGA X was used to generate the phylogenetic tree. Wheat proteins are colored according to their subfamily; proteins from other species are black. Rice subfamily names are used to identify subfamilies. The wheat proteins were given names based on a prior naming pattern. The bootstrap values are shown by dots on the branches: red dots, 76–100; yellow dots, 41–75; gray dots, ≥40. Ta, *T. aestivum;* At, *Arabidopsis thaliana;* Gm, *Glycine max;* Zm, *Zay maize;* Os, *Oryza sativa*.

Members of the BURP VI and BURP VII subfamilies all belong to three monocots (wheat, rice, and maize). The phylogeny of BURP genes followed species phylogeny almost exactly. For example, the PG1β-like subfamily could be assigned to two categories, monocotyledons and dicotyledonous BURP genes. The BNM2-like, BURP V (with five members found only in rice), and USP-like subfamilies contained only BURP members from the investigated dicots, which is consistent with previous results [6].

### Conserved motifs and gene structure of TaBURP

To better understand the diversity and similarity of wheat BURP domain proteins, we used the MEME software to identify motifs, finding 15 conserved motifs (Fig. 2). The results of these analyses suggest that the existing classification of BURP genes is appropriate. Within each subfamily, the motif numbers and their arrangement were almost identical. Specifically, the BURP VI subfamily contained ten motifs (10, 14, 1, 9, 12, 4, 3, 2, and 8), the BURP VII subfamily contained eight motifs (10, 9, 12, 6, 4, 3, 2, and 8), the BURP VIII subfamily contained six motifs (5, 9, 6, 4, 3, 2, and 8), the RD22-like subfamily contained nine motifs (10, 11, 9, 12, 6, 4, 3, 2, and 8), and the PG1β-like subfamily contained ten motifs (13, 7, 15, 9, 12, 6, 4, 3, 2, and 8).

**Fig. 2.**
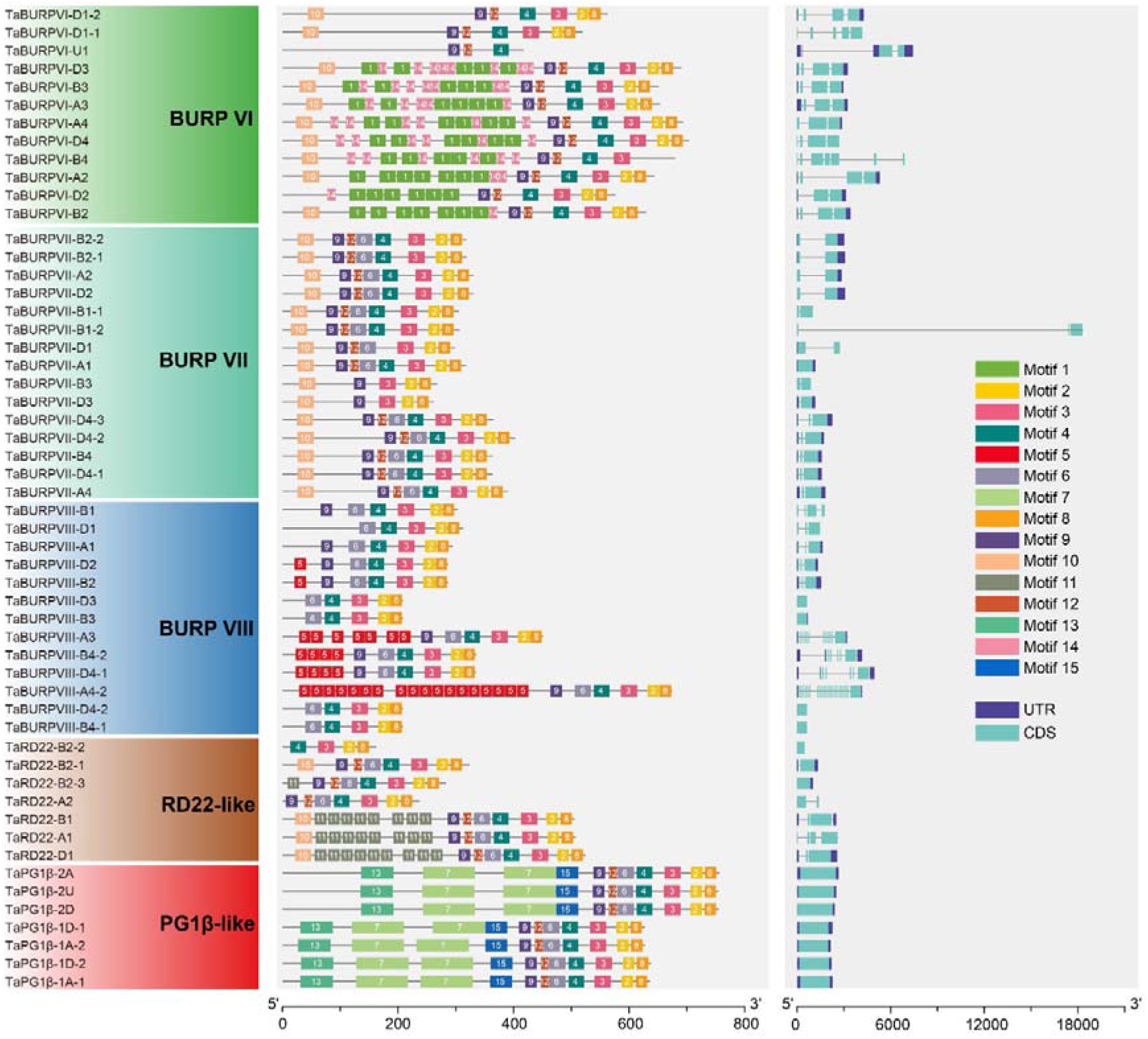
Motif and gene structure analysis of *TaBURP*s. Left: Maximum-likelihood (ML) phylogenetic tree deduced from the complete sequences of 54 *TaBURPs* (unrooted). Left: the background colors of subclades depict the five corresponding BURP subfamilies. Middle: the 15 motifs of BURP proteins identified by MEME. Different motifs are marked by distinct colored boxes and numbers. Right: gene structure of *TaBURPs*. UTR, CDS, and introns are indicated with purple boxes, green boxes, and black lines, respectively. Scale lines below the middle panel, the number of amino acids; Scale lines below the right panel, the number of amino acids bases.

Almost all wheat BURP members possessed motifs 4, 3, 2, 8 and 9 (**Additional file 2: Figure S1**). The exception was *TaBURP-U1*, which included only two motifs (12, 4). Generally, every subfamily contained one or more unique motifs (**Additional file 2: Figure S1**). For example, the PG1β-like subfamily contained three distinctive motifs: 13, 7, and 15. Motif 11 was exclusive to the RD22-like subfamily. The BURP VIII subfamily contained a unique and abundant motif 5, whereas the BURP VI subfamily comprised characteristic and substantial numbers of motif 1 and motif 14. This variation may be due to the loss of gene fragments during the evolutionary process.

### *Cis*-acting elements analysis of *TaBURPs* promoters

Many well-characterized stress-inducible promoter cis-elements confer enhanced plant tolerance to diverse abiotic stresses [29]. In an attempt to better comprehend and illuminate the potential regulatory functions of *TaBURP* genes under diverse stresses, we identified four types of cis-elements (abiotic stress response, light response, hormone response and growth-related cis-elements) in a 2000-bp promoter region upstream of the start codons (ATG) of the 54 *TaBURP* genes identified (Fig. 3 and **Additional file 1: Table S5**). All 54 *TaBURP* promoters had at least one stress-responsive element, and most *TaBURP* genes had light-response cis-elements and hormone-related (e.g., ABA and MeJA) cis-elements. These abundant cis-acting elements might help govern the spatial and temporal expression of genes, which in turn regulate plant growth and development as well as environmental stress response processes.

**Fig. 3.**
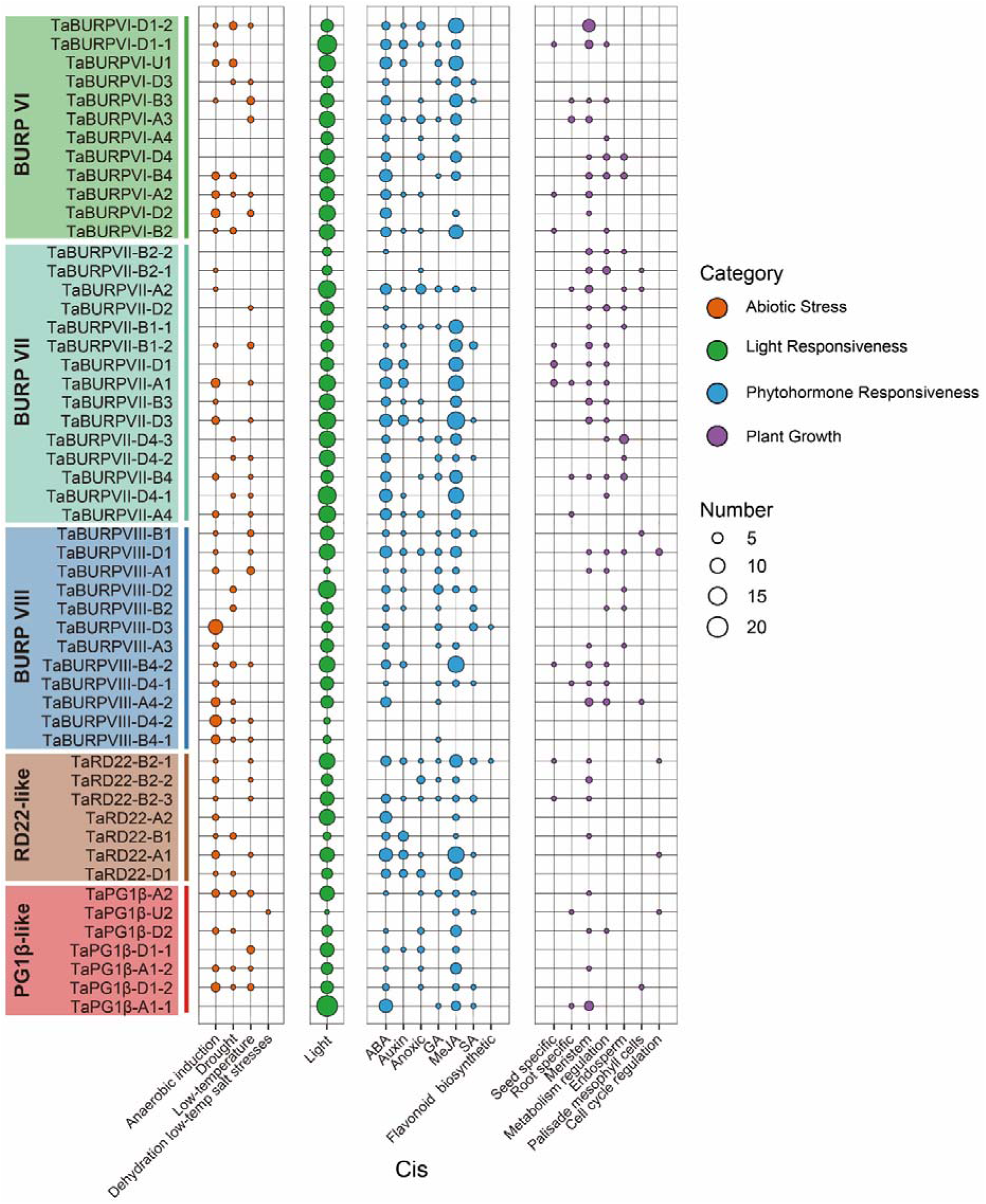
Distribution of the four types of cis-elements among the promoters of *TaBURP* genes. Circle color indicates the component type, while circle size indicates the number of components. *TaBURP* promoter cis-elements could be categorized into four types: abiotic stress response, light response, hormone response, and growth and development processes.

### Genomic location, gene collinearity, and duplication events of wheat BURP genes

We next investigated the chromosome distribution of the BURP gene wheat and its progenitors (*Ae. tauschii, T. urartu,* and *T. dicoccoides*). We found that twelve BURP genes of *T. urartu* were located on five chromosomes (3, 4, 5, 6, and 7); Twenty-three BURP genes of *Ae. tauschii* were located on six chromosomes (1D, 2D, 4D, 5D, 6D, and 7D); and thirty-two BURP genes of *T. dicoccoides* were located on thirteen chromosomes (1A, 1B, 2B, 3A, 3B, 4A, 4B, 5A, 5B, 6A, 6B, 7A, and 7B) (**Additional file 1: Tables S2**).

In wheat, the 54 TaBURP genes were unevenly distributed on 20 chromosomes, with no BURP genes on chromosomes 2A and 4B (Fig. 4a). Chromosome 6B contained the largest number of BURP members (7), whereas four chromosomes (1B, 2B, 2D, and 5A) each contained only one BURP gene. The distribution of *TaBURP* genes also differed among subgenomes, with subgenomes A, B, and D containing 14, 18, and 20 members, respectively (Fig. 4, **Additional file 1: Tables S1**). This heterogeneity between subgenomes may be the result of ancestral translocations and replication events.

**Fig. 4.**
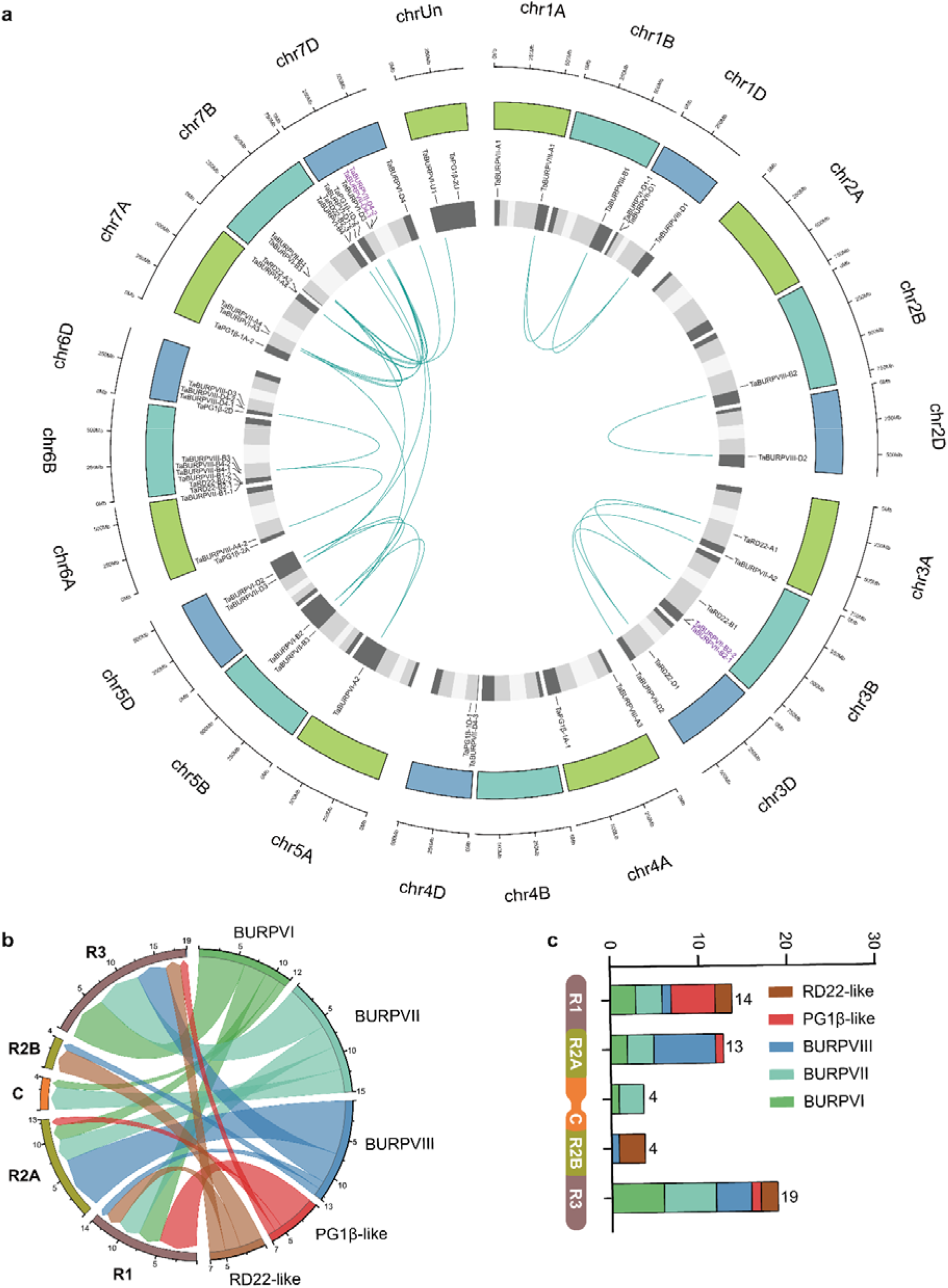
Location and gene duplication events of wheat BURP genes. **a** In a circular diagram, all BURP genes were mapped to their appropriate loci in the wheat genome. Subgenomes are shown in various hues of blue (outer track), whereas chromosomal segments are represented by shades of gray (inner track). Link lines (dark cyan) inside the circle represent gene pairs of segment duplication (24 pairs), whereas purple text represents gene pairs of tandem duplication (2 pairs). **b** The chord diagram indicates the number of BURP genes classified into different chromosomal positions. **c** A schematic diagram of the chromosomes depicting the diverse segments R1 and R3 (brown), R2A and R2B (light green), and C (orange). The number of genes from the different BURP subfamilies for each chromosome segment is shown in a bar plot.

Previous research have revealed that there are variances in the evolutionary rates of different regions of chromosomes. The rate of recombination events is generally considered to be lower near the central region of the chromosome (R2A, C, and R2B) and higher at the chromosome ends (R1 and R3) [30, 31]. Refinement of the locations of *TaBURP* genes showed that their chromosomal locations fell into two patterns (Fig 4b-c). Specificallly, 33 (61.1%) BURP genes were distributed in the distal region(R1 and R3), while 21 (38.9%) genes were located in the central region. Besides, genes from BURP VI, BURP VII, RD22-like, and PG1β-like subfamilies tended to be in distal chromosomal segments (R1 and R3), and collectivelly accounted for 51.9% of wheat BURP family genes (**Additional file 1: Table S6**). For the newly defined BURP VIII subfamily, eight members (14.8%) were situated in the central segment and five members (9.2%) were placed in the distal segment.

Segment duplication and tandem duplication are common duplication events that are instrumental in genome evolution and gene family formation [32]. In wheat, we identified two tandem duplications and twenty-four segment duplications, which combined accounted for 48.1% of BURP genes (**Fig. 4a and Additional file 1: Table S7**). This result suggests that duplication events may contribute to the expansion of the BURP gene in wheat. To analyze the selection pressure on BURP gene duplication pairs, we also analyzed the ratio of non-synonymous (Ka) to synonymous (Ks) substitutions. The results showed that all of the resulting Ka/Ks ratios were <1.0, implying that these gene pairs evolved mainly under purifying selection following gene duplication. Gene replication pairs may perform similar functions by purifying selective restriction differences (**Additional file 1: Table S7**).

Collinearity analysis can explain evolutionary relationships and polyploidy events. we conducted synteny analysis between wheat and two eudicots (Arabidopsis and soybean) and two monocots (rice and maize). Three wheat BURP genes are homologous to both rice and maize, but not to Arabidopsis and soybean, which is in line with the species phylogeny (Fig. 5a and **Additional file 1: Table S8**).

**Fig. 5.**
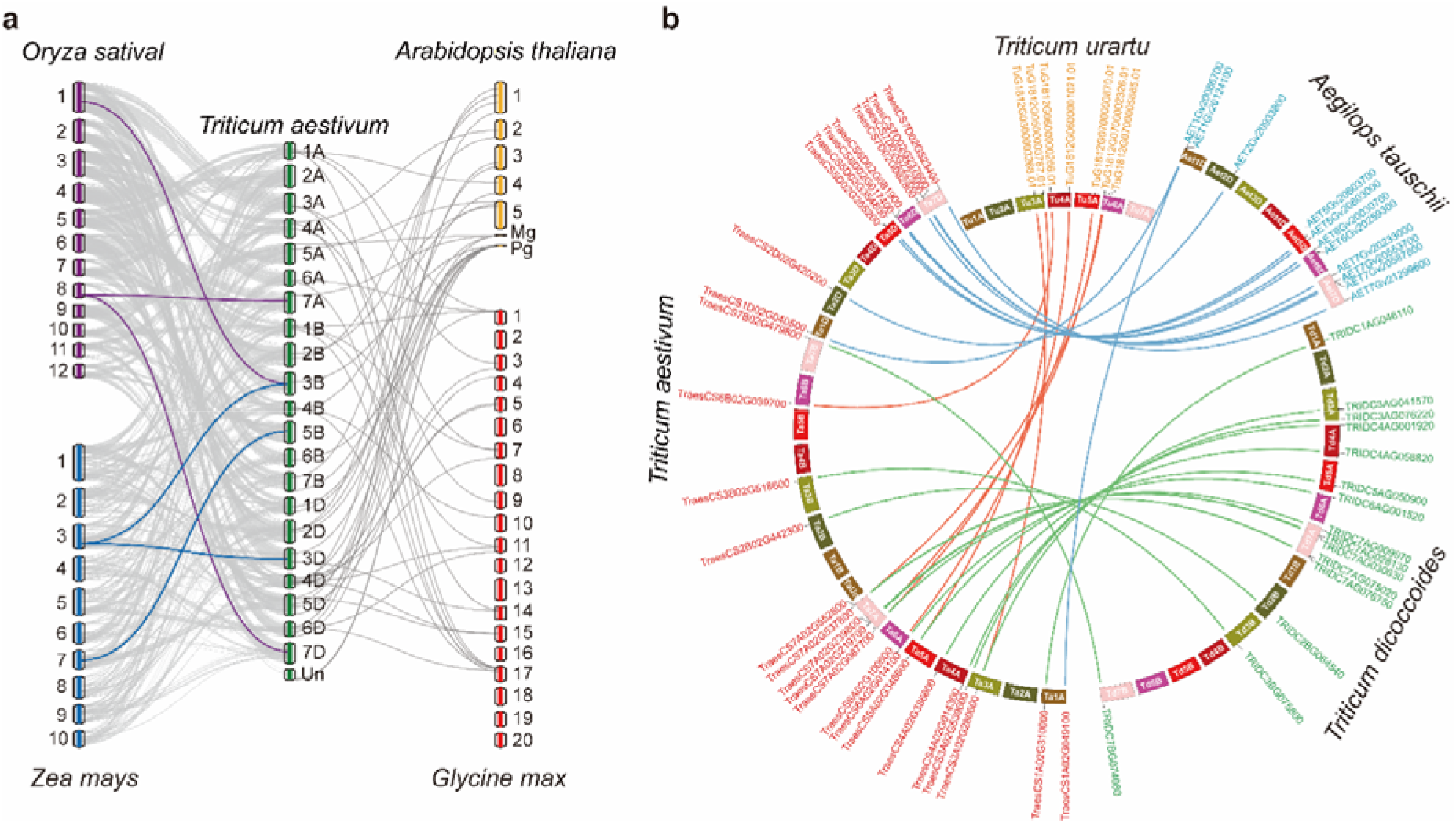
Synteny relationships of BURP genes with different plant species. **a** Yellow, red, green, purple, and blue rounded rectangles indicate chromosomes of Arabidopsis, soybean, wheat, rice, and maize, respectively. **b** Relationships between *Triticum aestivum* (AABBDD) and *Triticum urartu* (AA), *Aegilops tauschii* (DD), and *Triticum dicoccoides* (AABB), respectively. Red, yellow, blue, and green text on the oudside of the tracks represent BURP genes from *T. aestivum*, *T. urartu*, *Ae. tauschii*, and *T. dicoccoides*, respectively. Different-colored boxes denote different chromosomes, as indicated by white text inside boxes.

Subsequently, we performed synteny analysis between wheat and its ancestors. The results showed that 7, 11, and 15 pairs of orthologs between *T. aestivum* and its ancester: *T. urartu*, *Ae. tauschii*, and *T. dicoccoides*, respectively (Fig. 5b and **Additional file 1: Table S8**). After eliminating redundancy, totally 21 *TaBURPs* were derived from ancestral species, implying that these BURP members may have been formed by genome-wide duplication or polyploidization. The other BURP members from *T. urart*, *Ae. tauschii*, and *T. dicoccoides* (5, 12, and 17 members, respectively) were missed in *T. aestivum,* suggesting that they underwent genetic sequence losses during the generation of *T. aestivum*.

### Expression patterns of *TaBURPs* under ABA treatment and abiotic stresses

The BURP gene family is involved in multiple stress responses [5, 6, 23]. To explore the expression patterns of the *TaBURPs* genes in a variety of environmental stress conditions, we analyzed their expression profiles under drought, cold, salinity stress and as well as ABA treatment through RT-qPCR. Our results showed that many BURP genes are able to respond to a variety of stresses (Fig. 6). For example, fourteen *TaBURPs* were induced by all four treatments (red dots in Fig. 6); three genes (*RD22-B2-1, TaPG1β-D2, TaBURPVIII-B1*) were induced under cold, salt, and ABA treatment (light green dots); and two (*TaBURPVII-B2-1* and *TaBURPVII-A2*) were induced under drought, salt, and ABA treatments.

**Fig. 6.**
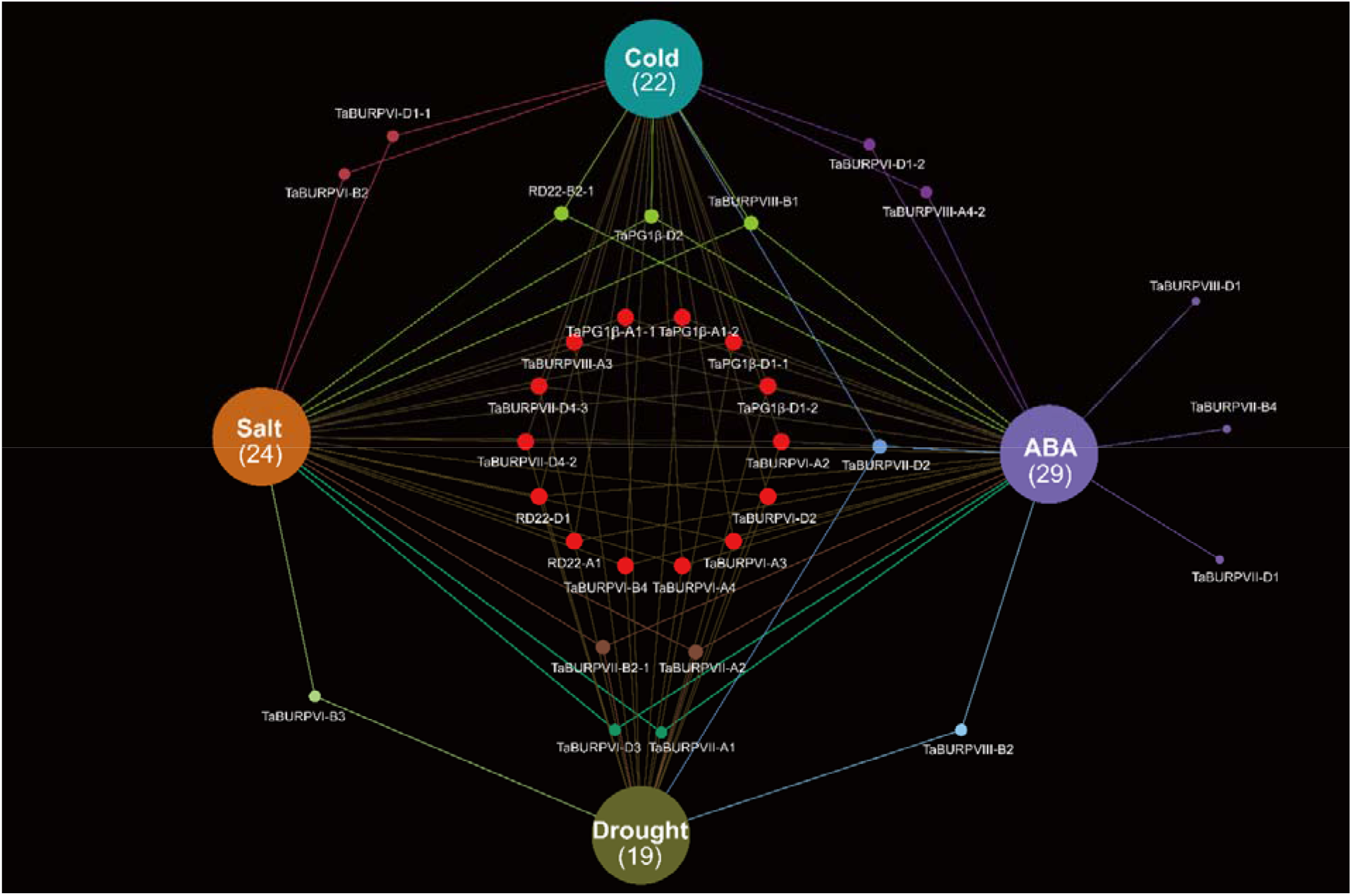
Wheat BURP gene responses to different stresses showed using a Venn network. Venn network plot generated using EVenn [33].

Nineteen *TaBURP* genes were expressed after drought treatment, showing four different expression trends. Six TaBURP genes showed a trend of increasing and then decreasing expression after simulated drought stress, with *TaBURPVII-B2-1*, *TaBURPVIII-A3* and *TaPG1β-D1-2* showing the most conspicuous (>10-fold) changes in expression (**Additional file 2: Fig. S2a**). In contrast, four genes had decreased expression following drought stress (**Additional file 2: Fig. S2b**). Six *TaBURP* genes showed a distinct pattern in which their expression decreased, increased, and then decreased again following drought stress (**Additional file 2: Fig. S2c**). However, the other three genes showed no clear pattern (**Additional file 2: Fig. S2d**).

Under salt stress, a total of 24 TaBURP genes were expressed and showed three expression patterns. Briefly, four *TaBURPs* experienced rapid increases in expression in response to high salinity (**Additional file 2: Fig. S3a**). Similarly, six *TaBURP* genes showed increased expression by 5 h after the start of salinity stress, in most cases peacking after 8 h (**Additional file 2: Fig. S3b**). Among these, the expression of *TaBURPVI-B3, BURPVII-B2-1*, *BURPVIII-A3*, and *TaPG1β-D1-1* was sharply increased after 8 h of salinity stress (>20-fold), indicating their potential importance in plant osmotic stress tolerance. Six *TaBURP* genes had decreased expression after salinity stress (**Additional file 2: Fig. S3c**), implying that they might involved in salt stress responses.

Following cold stress treatment, the expression levels of three Ta*BURP* genes increased and then returned to their initial levels (**Additional file 2: Fig. S4a**). For 17 *TaBURP* genes, expression levels first increased slightly, then increased more rapidly, and subsequently decreased following cold stress (**Additional file 2: Fig. S4b**). These genes reached their maximum expression after 8 h, except for *TaBURPVI-A2* and *TaBURPVI-B4*. The expression of *TaBURPVII-A1* and *TaBURPVII-D1* initially decreased rapidly under cold stress but then returned to normal levels (Additional file 2: Fig. S7c). Three genes (*TaBURPVII-D4-2, TaBURPVII-D4-3,* and *TaBURPVIII-A3*) showed fluctuating expression patterns under cold stress (**Additionalfile2: Fig. S4d**).

ABA treatment induced the expression of 29 *TaBURP* genes. Among these, the expression of 21 BURP genes peaked at 8 h but other 8 genes were hardly detectable at 8 h after ABA treatment (**Additional file2: Fig. S5**). Overall, these data show that different abiotic stressors have varied and extensive effects on the expression of wheat BURP genes.

### Identification and annotation of potential BURP target genes

Next, we generated a protein–protein association network was examined using the STRING online server (https://string-db.org/cgi). The interaction network contained 21 wheat BURP proteins and 20 target proteins (Fig. 7).

**Fig. 7.**
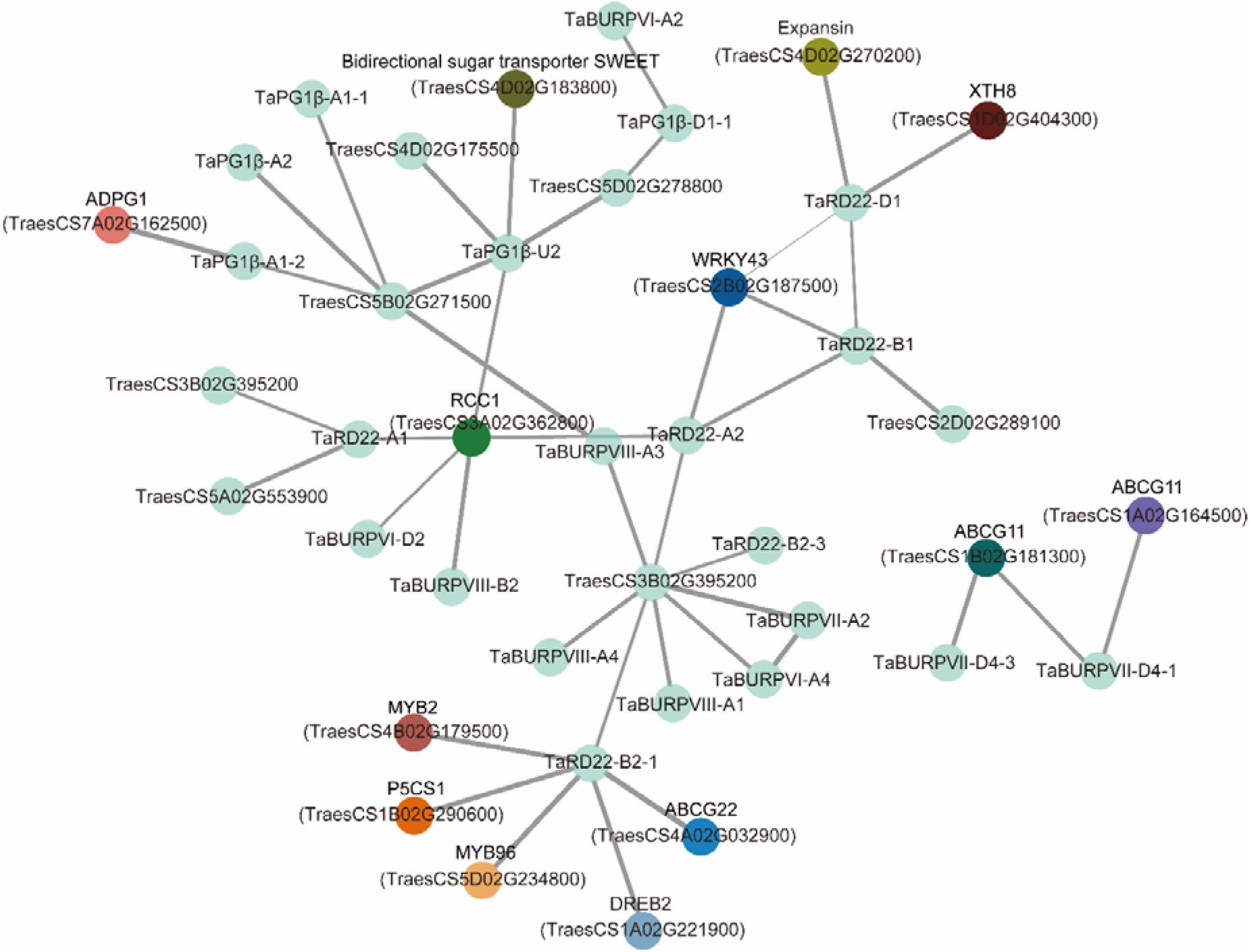
Putative interaction network of BURP proteins in wheat. Colors other than light green represent annotated genes. RCC1, regulator of chromosome condensation; XTH8, xyloglucan endotransglucosylase/hydrolase; ABCG22, ABC transporter G family member 22; ABCG11, ABC transporter G family member 11; DREB2, dehydration-responsive element-binding protein 2; P5CS1, delta1-pyrroline-5-carboxylate synthase 1; ADPG1, Arabidopsis dehiscence zone polygalacturonase1. The thickness of the gray line indicates the confidence score from the STRING website.

TaRD22-B2-1 can interact with six wheat proteins (TraesCS4B02G179500, TraesCS1B02G290600, TraesCS5D02G234800, TraesCS1A02G221900, TraesCS4A02G032900 and TraesCS3B02G395200), which were involved in responses to diverse abiotic stresses (e.g., DREB, MYB, P5CS and ABCG) [34–39]. Among TaRD22-A1/A2, TaPG1β-U2, TaBURPVI-D2 and TaBURPVIII-B2 can interact with RCC1 protein (TraesCS3A02G362800), which RCC1 protein (TCF1) can enhance tolerance for chilling and freezing by regulating lignin biosynthesis [40]. TaBURPVII-D4-1/3 can interact with ABCG11, the *ABCG11* gene is required for epidermal lipid secretion [41]. Meanwhile, TaRD22-D1 can interact with TaRD22-B1 and three other wheat proteins (TraesCS4D02G270200, TraesCS1D02G404300 and TraesCS2B02G187500), which were WRKY43, XTH8 and Expansin, WRKY transcription factors (TFs) play an important role of developments and stresses; XTH8 is essential for loosening cell walls and dissolving the cellulose–xyloglucan matrix [42, 43]; Expansins have many important biological functions, such as seed development, fruit ripening and stress tolerance [44]. TaPG1β-A1-2 can interact with ADPG1 protein (TraesCS7A02G162500) and TraesCS5B02G271500, ADPG1 is essential for anther dehiscence through degradation of pectin [45]. We also predicted that TaPG1β-U2 interact with SWEET (TraesCS4D02G183800), RCC1 (TraesCS3A02G362800) and three other wheat proteins (TraesCS4D02G175500, TraesCS5D02G278800 and TraesCS5B02G271500), SWEET family members participate in important physiological processes such as phloem loading, pollen development, nectar secretion, seed germination, leaf senescence, and fruit development, and play a key role in host–pathogen interactions and various responses [46–54]. These results provided valuable information for the further functional characterization of *TaBURP* genes.

### Tissue-specific expression patterns of *TaBURP* genes

To investigate the temporal and spatial expression patterns of wheat BURP genes across different tissues at distinct developmental stages, we using available expression data from WheatOmics 1.0 (http://202.194.139.32/) [55]. *TaBURP* expression patterns varied greatly across tissues and development stages (Fig. 8), with large variations both among and within subfamilies. For example, BURP VI subfamily members were predominantly expressed in roots, whereas RD22 subfamily members were not expressed in roots. Three RD22-like subfamily members (*TaRD22-A1, TaRD22-B1*, and *TaRD22-D1*) were highly expressed in stems, leaves, spikes, and seeds. Half of the BURP VIII subfamily genes were not expressed in any tissues except spikes and grains. More than half of PG1β-like subfamily BURP genes were expressed in multiple tissues, with the remainder expressed only in spikes. As seen in the Fig. 8, 30 BURP genes (55.6%) were expressed in spikes, among which 10 (*TaBURPVII-D3*, *TaBURPVII-A4, TaBURPVII-B4, TaBURPVII-D4-1, TaBURPVIII-B2, TaBURPVIII-D2, TaBURPVII-B3, TaPG1β-A2, TaPG1β-D2,* and *TaPG1β-U2*) were spike-specific. Expression profiles showed that *TaBURP* genes may play distinct roles during plant growth and development.

**Fig. 8.**
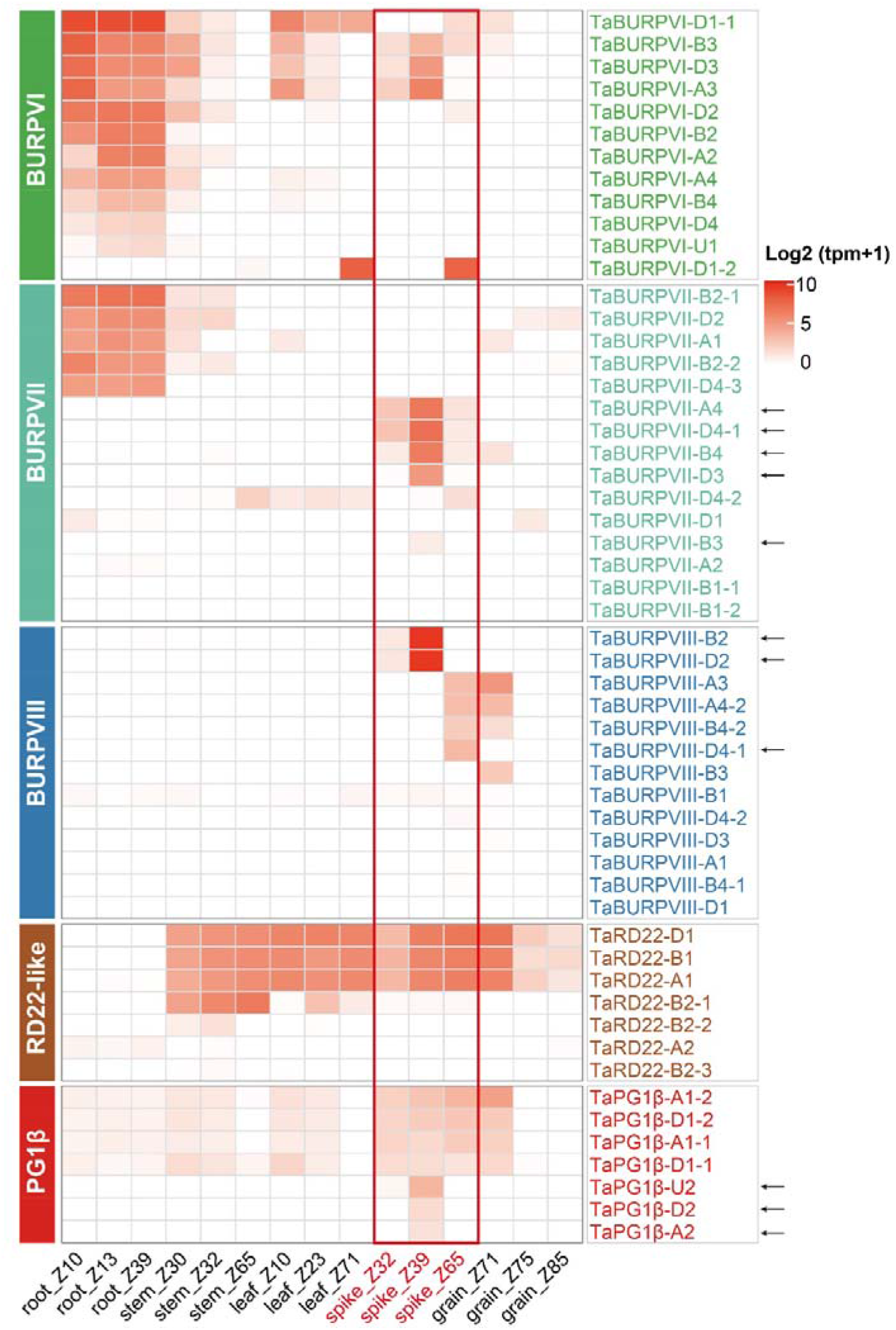
Heat map of the transcript levels of 54 *TaBURP* genes across tissues at different developmental stages. The red rectangle indicates expression patterns of *TaBURP* genes in the spike. Black arrows indicate genes specifically expressed in the spike.

## Discussion

### Overview of the wheat BURP gene family

BURP genes have been systematically identified and characterized in a variety of plants, including Arabidopsis, rice, soybean, and cotton. These genes are involved in all aspects of plant growth and development, such as seed, root, spike, and embryo development. They also play an important role in diverse stress responses. Therefore, they are promising candidates for plant genetic improvement and crop breeding.

Recent advances in sequencing technology have accelerated the assembly of the wheat genome, resulting in new, high-quality genomic data that allowed the accurate identification of the BURP gene family [56]. In this study, we identified 54, 12, 23, and 32 putative BURP genes in *T. aestivum*, *T. urartu, Ae. tauschii,* and *T. dicoccoides,* respectively. Of the 54 wheat BURP genes, 21 showed collinearity with their ancestral species based on synteny analyses, indicating that whole-genome duplication or polyploidization was an important factor driving *TaBURP* expansion. However, several homologs of *T. urartu, Ae. tauschii,* and *T. dicoccoides* BURP genes were missing in *T. aestivum,* indicating that chromosomal segment deletion or rearrangement, point mutations, and gene insertions or deletions all occurred during the evolution of *T. aestivum* [6, 57].

Phylogenetic analysis revealed that the wheat BURP gene family can be divided into five subfamilies (BURP VI, BURP VII, BURP VIII, RD22-like, and PG1β-like). Our results differ slightly from those of previous studies due to the approach we used to construct the phylogenetic tree and the species used. BURP genes from monocots and dicots were found in the RD22-like and PG1β-like subfamilies. Within each subfamily, the homologs of dicot and monocot BURP genes were divided into two lineages, showing that they evolved before monocots and dicots diverged [9]. These findings are consistent with previous research [6, 9]. However, BURP genes from monocots (BURP V, BURP VI, BURP VII, and BURP VIII) and from dicots (USP-like, BNM-like) clustered separately, suggesting that they may have evolved separately after the monocot-dicot split [5, 7, 58].

### Subfamily-specific amplification might facilitate high adaptation in wheat

The hexaploid nature of wheat and the substantial size of the BURP gene family provide ideal opportunities to study gene duplication and the evolutionary fate of genes following polyploidization. With 54 genes, wheat has one of the highest numbers of BURP genes among the flowering plants. BURP genes are 3.17 times more abundant in wheat than in rice (17) (Additional file 2: Table S1), probably as a result of the hexaploidy of wheat. In particular, the BURP VI (12), BURP VII (15), and BURP VIII (13) subfamilies contained 4-, 7.5-, and 13-fold more genes in wheat than in rice, respectively. It is likely that these BURP families expanded through gene duplication events after lineage differentiation.

The rate of gene evolution is generally higher in the distal regions of chromosomes than in the proximal regions. Numerous rapidly evolving genes are situated in evolutionary hotspots, such as distal telomeres, traditionally thought to be targets for recombination [30, 31]. Recently, genome-wide sequencing of wheat revealed that genes associated with stress responses and external stimuli tend to be located in distal chromosomal regions, whereas those pertinent to transcription and translation regulation, apoptosis and signal transduction, and cell cycle regulation are abundant in proximal chromosomal segments [56, 59]. That is, genes that are designed to adapt quickly to various circumstances are generally located in the terminal regions of chromosomes, while those that are important and conserved tend to be in central segments of chromosomes. In our study, the expanded subfamilies tended to be in the distal chromosome segments. Specifically, 75%, 60%, 85.7%, and 57.1% of the BURP VI, BURP VII, PG1β-like and RD22-like genes, respectively, were located in the distal chromosome segments (e.g., R1 and R3) (Additional file 1: Table S6). The genes of these four subfamilies tended to be anchored in distal telomeric regions, suggesting that they have higher evolved ratios. The higher incidence of duplication events in distal telomeric segments is plausibly responsible for the expansion of these subfamilies, which in turn could benefit from rapid adaptation to diverse environmental conditions [28].

However, the BURP VIII subfamily was an exception, with 62% of its genes located in proximal telomeric segments, suggesting that these genes may be highly conserved. The BURP VIII subfamily with highly conserved functions may have evolutionary advantages due to its gene redundancy, which could effectively reduce the adverse effects of mutations. Overall, the expansion of wheat BURP genes may allow for rapid adaptation to various environmental conditions, thus contributing to its global distribution.

### Putative functions of *TaBURP* genes in wheat development

Spatiotemporal expression patterns of genes are generally recognized as an important means of studying their function [60]. We identified the BURP VI subfamily as a monocotyledon-only subfamily, which is consistent with earlier findings. Most members of the BURP VI subfamily were strongly expressed in roots. Roots are vital organs for nutrient absorption. After phosphorus deficiency treatment, TaBURPVI-D3, TaBURPVI-A2, TaBURPVI-B3, TaBURPVI-B4, TaBURPVI-D1-1, TaBURPVI-D2, TaBURPVI-D4, and TaBURPVI-A3 showed increased expression [61]. This suggests that several wheat BURP VI subfamily members are involved in phosphorus absorption. In addition, *TaBURPVII-A2*, *TaBURPVII-B2-1, TaBURPVII-B2-2,* and *TaBURPVII-D2* were mainly expressed in roots, suggesting that these members are involved in root function.

Many BURP genes may be involved in the development of reproductive organs. For example, the three putative wheat homologs of *OsBURP4 (BURPVIII-B2/D2/D4-1*) had expression restricted to the spike, suggesting that they are pertinent to spike development. OsBURP13 is also known as OsRAFTIN and belongs to the rice BURP VII subfamily. This protein is necessary for pollen formation [18]. BURPVII-A4 (TaRAFTIN1a) and BURPVII-B4 (TaRAFTIN1b), two wheat orthologs of OsBURP13, accumulate in Ubisch bodies and are required for pollen formation [18]. *OsBURP15 (RA8*) is an anther-specific gene [19]. In this study, six wheat orthologs of *OsBURP15 (BURPVII-B3, BURPVII-D3, BURPVII-A4, BURPVII-B4, BURPVII-D4-1*, and *BURPVII-D4-2*) were expressed primarily in spikes, implying that they are important in spike development.

Wheat BURP members might be indispensable for seed development. Overexpression of cotton RD22-like proteins (GhRDL1) increases cotton seed mass and biomass [62]. We found that three wheat RD22-like proteins (*TaRD22-A1*, *TaRD22-B1*, and *TaRD22-D1*) have high levels of expression during seed development, indicating that they are important in seed development. AtPGL3 can regulate pectin solubilization and degradation. In our study, the *AtPG2 (AtPGL3*) homologs *TaPG1β-A1, TaPG1β-B1,* and *TaPG1β-U1* were preferentially expressed in the anther, indicating that they may function in pollen development, possibly by degrading pectin in the pollen cell wall [21, 63]. Moreover, several RD22-like family members can control cell wall lignin content. For example, ectopic expression of *GmRD22* in Arabidopsis and rice resulted in increased cell wall lignin content under salinity stress [64].

### Putative functions of *TaBURP* genes in wheat stress tolerance

Cis-elements are essential for transcriptional regulation and expression. We found that wheat BURP promoters contained a variety of cis-acting elements, including those related to abiotic stress response, light response, hormone response, and growth. Abiotic stressors are the primary constraint on plant development and distribution. According to our cis-element analysis, 48 *TaBURP* genes shared more than one abiotic stress response element. For example, under drought, salt, and ABA treatments, two homologous *RD22* genes, *TaRD22-A1* and *TaRD22-D1*, showed markedly and rapidly elevated expression, providing compelling evidence for their participation in stress tolerance. Furthermore, overexpression of *OsBURP16,* a member of the rice PG1-like subfamily, has previously been recommended to enhance rice sensitivity to drought, salinity, and cold stress [65]. Drought, salinity, cold, and ABA treatment activated four wheat orthologs of *OsBURP16 (TaPG1β-A1-1, TaPG1β-A1-2, TaPG1β-D1-1,* and *TaPG1β-D1-2),* indicating that they have similar activities in plant stress tolerance.

The ABA signaling pathway is also thought to be associated with stress response [66]. In this study, ABA treatment induced the expression of 29 *TaBURP* genes at various time points. Among these, 28 contained at least one ABA-responsive cis-element, implying that these genes are involved in the ABA signaling pathway via ABA-responsive cis-acting elements. These results indicate that TaBURPs are heavily involved in the response to a variety of stressors, suggesting that they may have conferred wheat adaptive plasticity and thus helped global dispersion.

## Methods

### Identification of BURP genes in wheat and its progenitor

All nonredundant BURP protein sequences of Arabidopsis, soybean, rice, maize, *Ae. tauschii, T. dicoccoides,* and *T. aestivum* were downloaded from Ensembl Plants, while those of *T. urartu* was obtained from the MBKBase website (http://www.mbkbase.org/Tu/).

A Hidden Markov Model (HMM) profile of the BURP domain (PF03181) was downloaded from the Protein family database (Pfam) [66]. Then, the HMMER 3.2.1 was used to identify the BURP genes for *T. aestivum, Ae. tauschii, T. urartu,* and *T. dicoccoides* [67]. In parallel, a local BLASTP program was used to screen putative BURP members for *T. aestivum* and its progenitor species, with an *E*-value cut-off of <1e^-10^. To improve the accuracy of the search results, only those candidates with an identity value >75% were used for subsequent analysis. Results from HMMER and BLASTP were mixed and validated on Pfam and SMART (http://smart.embl.de/smart/batch.pl) databases.

Altogether, we identified 54, 12, 23 and 32 BURP genes in *T. aestivum, T. urartu, Ae. tauschii,* and *T. dicoccoides,* respectively. Subsequently, to explore the protein properties of all identified BURP genes, the MW, pI, and GRAVY of each BURP protein were predicted using ExPASY (http:web.expasy.org/protgram/) [68]. BURP protein subcellular location prediction was performed using WoLF PSORT (https://wolfpsort.hgc.jp/).

### Multiple sequence alignments and phylogenetic analysis

All identified BURP protein sequences from Arabidopsis, soybean, rice, and maize, as well as the four canonical BURP domain proteins (BNM2, VfUSPs, RD22, LePG1β), were merged and aligned using MUSCLE [69]. The alignment results were then used to deduce a phylogenetic tree using MEGA X with the maximum likelihood method [70]. The substitution model was the JTT model with 1000 bootstrap replications, according to the Bayesian information criterion. The resulting tree file was then visualized with Evolview V2 [71].

### Chromosomal location, gene duplication, and synteny analysis of BURP genes

The distribution of BURP genes was determined by the physical localization of the genomic database of wheat and its progenitor. Gene duplication events were determined according to previous research [32]. A synteny analysis was performed using MCScan X [72]. The synteny analysis results were visualized with Circos software [73].

The selection pressure of each BURP gene duplication event was calculated as the Ka/Ks ratios using Kaks calculator 2.0 [74]. In general, Ka/Ks < 1 indicates purifying selection, Ka/Ks = 1 signifies neutral selection, and Ka/Ks > 1 indicates positive selection. The dispersal times of these gene replication pairs were inferred from earlier studies. The synteny relationships of BURP genes between wheat and its ancestral species were analyzed. The syntenic relationships between rice, maize, Arabidopsis, and soybean were also investigated.

### Interaction analysis of TaBURPs and their targets

A BURP protein interaction network map of different species (rice, maize, Arabidopsis, soybean, and wheat) was constructed using STRING (https://cn.string-db.org/) [75] to identify wheat BURP protein functions. The proteins in the interaction network were then aligned to the wheat genome, retaining the wheat genes with the highest BLAST scores or with annotated functions. Subsequently, all retained genes were used to construct an integrated interaction network using Cytoscape v3.8.0 [76].

### Analysis of structure, motifs, and cis-elements of BURP genes

To understand the exon-intron organization of *TaBURP* genes, the gene sequences of *TaBURP* genes were aligned and visualized with the corresponding genome-wide sequence via TBtools [77]. The conserved motifs of BURP proteins were identified using MEME [78] with the following settings: the size distribution was 0 or 1 occurrence per sequence; the motif count was 15; and the motif width ranged from 6 to 100 residues. Then, 2-kb sequences from promoter regions of *TaBURP* genes were used to predict regulatory elements via the PlantCARE database (http://bioinformatics.psb.ugent.be/webtools/plantcare/html/) [79]. With the prediction results from this site, a matrix bubble plot was generated using the ggplot2 package [80].

### Gene expression analysis

Gene expression data were downloaded from WheatOmics 1.0 (http://202.194.139.32/) as transcripts per million [55]. These transcriptome datasets contained expression data for five high-level tissues (root, stem, leaf, spike, and grain) at ten time-point stages (Z10, Z13, Z23, Z30, Z32, Z39, Z65, Z71, Z75, and Z85) [81]. Subsequently, expression levels of BURP genes were visualized using heatmaps with the ComplexHeatmap package [82]. A clustering analysis was performed within each subfamily based on their expression profiled (metric, Euclidean; method, Average).

## List of abbreviations

AA: amino acid
ABA: abscisic acid
BURP: BNM2, USP, RD22, and PG1β
BNM2: a microspore protein
Dyad: the dyad stage
GA: gibberellins
HMM: hidden Markov model
MeJA: methyl jasmonate
MMC: the microspore mother cell stage
MP: the young microspore stage
MW: molecular weight
Ka: nonsynonymous substitution rate
Ks: synonymous substitution rate
PG1β: a non-catalytic β-subunit of the polygalacturonase isozyme
pI: isoelectric point
RD22: a dehydration-responsive protein
SA: salicylic acid
USP: an unknown seed protein

## Availability of data and materials

All data generated or analyzed during this study are included in this published article and its supplementary information files.

## Acknowledgments

We thank the National Medium Rice Genebank at the China National Rice Research Institute and the International Wheat Genome Sequencing Consortium (IWGSC) for online data support.

